# Introductory gestures before songbird vocal displays are shaped by learning and biological predispositions

**DOI:** 10.1101/2020.03.28.013052

**Authors:** Shikha Kalra, Vishruta Yawatkar, Logan S James, Jon T Sakata, Raghav Rajan

## Abstract

Introductory gestures are present at the beginning of many animal displays. For example, lizards start their head-bobbing displays with introductory push-ups and animal vocal displays begin with introductory notes. Songbirds also begin their vocal displays by repeating introductory notes (INs) before producing their learned song and these INs are thought to reflect motor preparation. Between individuals of a given species, the acoustic structure of INs and the number of times INs are repeated before song varies considerably. While similar variation in songs between individuals is known to be a result of learning, whether INs are also learned remains poorly understood. Here, using natural and experimental tutoring with male zebra finches, we show that mean IN number and IN acoustic structure are learned from a tutor, independent of song learning. We also reveal biological predispositions in IN production; birds artificially tutored with songs lacking INs still repeated a short-duration syllable, thrice on average, before their songs. Overall, these results show that INs, just like elements in song, are shaped both by learning and biological predispositions and suggest multiple, independent, learning processes underlying the acquisition of complex vocal displays.

## INTRODUCTION

Animal produce various complex displays to communicate with their conspecifics [1]. Many of these communicative displays begin with the repetition of introductory gestures. For example, Anolis lizards produce a few introductory “push-ups” or “tail-flicks” before starting their head bobbing display [2,3], frogs and toadfish produce introductory vocalizations before their advertisement calls [4,5] and a number of songbirds also produce introductory vocalizations before the start of their complex songs [6–17]. Many different functions have been attributed to these introductory gestures, including, an “alerting” function [5,18,19], a species-specific signal that aids learning of song [20], a “local-dialect” signal [21] and more recently, a role in motor “preparation” [22,23]. Given that these introductory gestures are a part of both learned and unlearned displays, the extent to which they are learned, in the context of learned displays like bird song, remains poorly understood.

Bird song is a well-studied example of a complex vocal display that is learned by imitation from a tutor [24–26]. Many different species of songbirds, including the zebra finch, begin their displays with the repetition of a short, simple, vocalization called an introductory note (IN) [6–12,14,16] before producing their more complex song. Among individuals of a given species, both the repetition and acoustic structure of INs vary considerably [23,27–29]. What is the source of this variation? Variation in elements of song between individuals is a consequence of learning [8,10,24,25]. Similarly, variation in IN number and IN acoustic structure could also be learned from a tutor. Alternatively, as predicted by the motor preparation hypothesis [22,23], variation in IN number and structure across birds could be correlated with variation in their respective songs. Finally, variation in IN number and structure across birds could also be a result of biological predispositions similar to the biological predispositions in the production of elements of song [30]. Here, we examine these different predictions in the zebra finch, a well-studied songbird [14].

Adult male zebra finches also begin their vocal displays by repeating an IN before their song (Fig. 1A) [8,10,14]. Both mean IN number and IN acoustic structure vary considerably between birds (Fig. 1A, 1B) [23,28,31]. Here, using natural and experimental tutoring methods, we show that the mean number of IN repeats before song and IN acoustic structure are learned from a tutor and this IN learning is independent of song learning. Second, by experimentally tutoring birds with songs lacking INs, we reveal the presence of biological predispositions in IN production; such experimentally tutored birds still repeated a short-duration vocalization, thrice on average, before their songs. Overall, our results demonstrate that, similar to song elements, INs are shaped by both learning and biological predispositions and suggest that the acquisition of complex vocal displays involves independent acquisition of INs and song.

**FIGURE 1.**
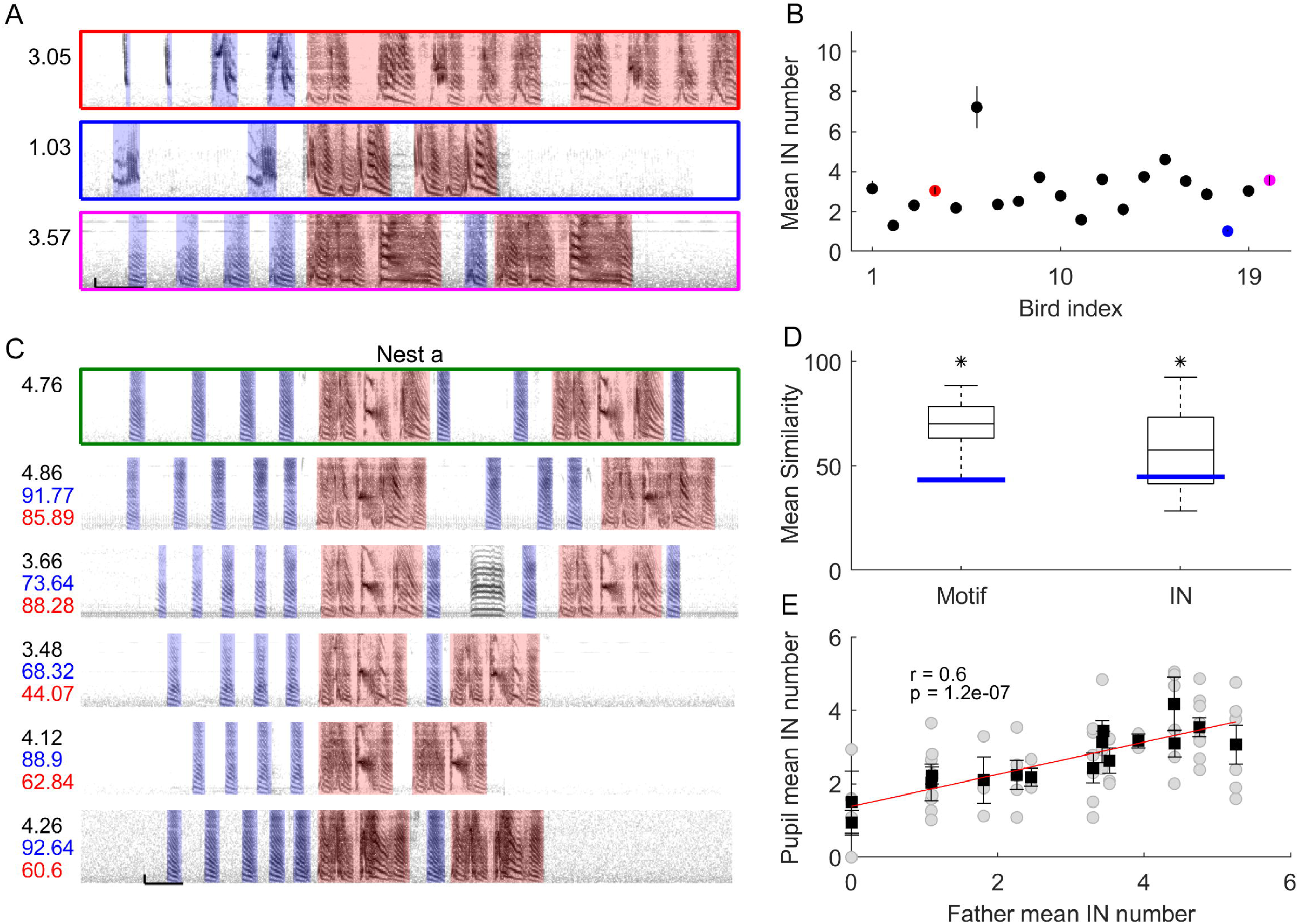
Mean IN number and IN acoustic structure are correlated between fathers and their normally reared sons. (A) Spectrograms of song bouts of 3 different zebra finches. Red shading highlights song motif and blue shading highlights INs. (B) Mean IN number varies considerably across 20 unrelated birds. Circles and whiskers represent mean and s.e.m for individual birds. Colours represent birds from (A). (C) Example spectrograms of one song bout of a father (green box) and 5 sons showing introductory notes (blue shaded boxes) and common motifs for each bird (red shaded boxes). Black numbers on the left indicate mean IN number for that bird. For sons, blue and red numbers represent mean similarity to father’s INs and motif respectively. (D) Mean song similarity and mean IN similarity between fathers and sons. Blue line represent chance level similarity (mean similarity between father from one nest and pupils from other nests). * represents p < 0.05, Wilcoxon rank sum test (E) Mean IN number of father vs. mean IN number of son. Grey circles represent individual birds, black squares and whiskers represent mean and s.e.m. for individual nests. Red line represents regression line.

## MATERIALS AND METHODS

All procedures at IISER Pune were conducted after approval from the Institute Animal Ethics Committee (IAEC), IISER Pune and were in accordance with CPCSEA (Committee for the Purpose of Control and Supervision of Experiments on Animals), New Delhi. All procedures at McGill were conducted following approval from the McGill University Animal Care and Use Committee in accordance with the guidelines of the Canadian Council on Animal Care.

Zebra finches obtained from an outside vendor or bred in IISER Pune or McGill were used for all experiments. Detailed methods are present in Supplemental Information.

### Song recordings

Songs were recorded in custom-made sound attenuation boxes (NewTech Engineering Systems, Bangalore, India; TRA Acoustics, Ontario, Canada) by placing an omnidirectional microphone (AKG 517; Countryman Associates, Menlo Park, CA) on top of the bird’s cage. Signals from the microphone were band-pass filtered, digitized and recorded to a computer (n=132 birds). All birds were recorded when they were adults (> 85 days post-hatch) and only undirected songs in the absence of any other bird were recorded. For all analyses, only files with > 1.9s of silence before and after song bouts in a file were considered.

### Experimental groups

We used 4 different experimental groups in our study.

#### Normally reared birds

All birds used in this group (n = 65, nests =16) were bred at IISER, Pune. A total of 16 nests were analysed with 13 nests having atleast 3 offspring (range: 1-9). Juveniles were housed with their parents and siblings until they were 50-94 days old, after which they were transferred to the colony and housed with other males from same or different families.

For all other groups, we used naive birds (i.e. birds that had not previously learned song) to test various hypotheses about IN learning. In order to prevent song learning from their father, juvenile zebra finches were separated from their father around phd 10 (range phd 6-16) and reared with their mother and siblings until phd 35 (range: phd 34-54). Previous studies have shown that exposure to the father’s song before phd 25 does not significantly impact learning [32], so our birds are unlikely to have learned their father’s song during this period. Once juveniles had reached nutritional independence they were separated and housed individually in small cages (except for 2 birds who were housed together). These birds were then used for the different groups as outlined below.

#### Playback tutored birds

Using active tutoring methods [33,34], birds were tutored with synthesized zebra finch songs that were played back through a speaker (Fig. 5A; n=22 birds from 11 different nests) at McGill (n=13) or at IISER Pune (n=9).

#### Socially tutored birds

14 birds were tutored with an adult male different from their father using a social tutoring paradigm [34,35] (Fig. 2A; n=10 at IISER Pune and n=4 at McGill; total n=14 birds from 11 different nests). For tutoring, a cage with an adult male (tutor) was placed next to the juvenile’s cage. For the 10 birds tutored at IISER Pune, we chose tutors with a different mean IN number than the father (mean difference in IN number between father and tutor – 2.75; range: 1.37 – 3.48). Birds tutored at McGill were part of a different study where song (but not IN) learning was described [35]. For these birds, INs of the tutors was not a consideration during the choice of tutor. Therefore, to investigate IN learning, only a subset of these tutored birds were analyzed here. Specifically, only socially tutored birds in which tutors produced a different number of INs than the biological father (mean difference in IN number between father and tutor – 2.06; range: 0.5-4.78) were included in these analyses.

**FIGURE 2.**
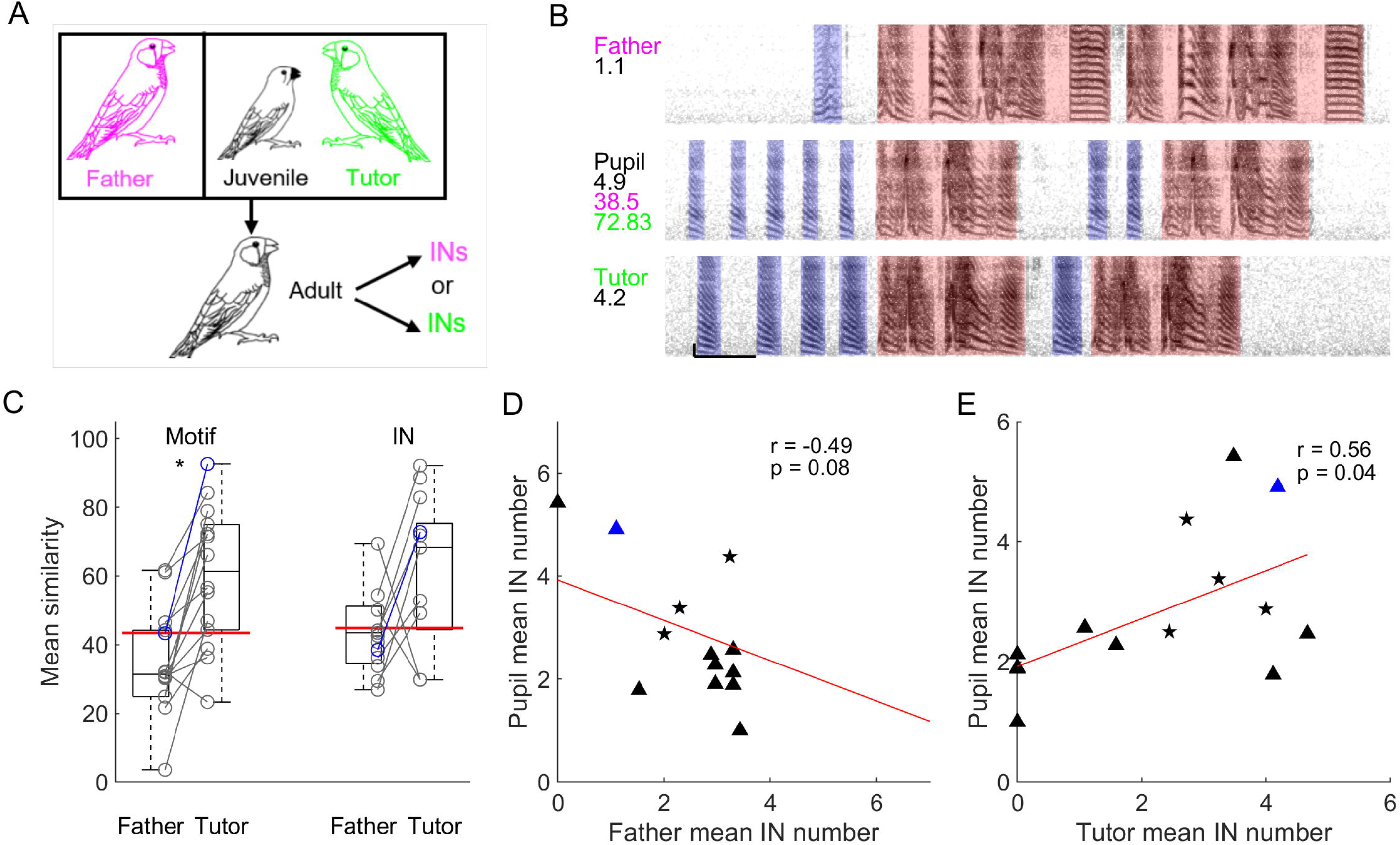
Mean IN number and IN acoustic structure, for socially tutored birds, is correlated with social tutor, not father. (A) Schematic of social tutoring paradigm. (B) Example spectrograms of one song bout of father (top), pupil (middle) and social tutor (bottom) showing introductory notes (blue shaded boxes) and common motifs for each bird (red shaded boxes). Black numbers on the top left side of each spectrogram represent mean IN number for that bird. For the pupil, magenta and green numbers on the left represent mean similarity to father’s and social tutor’s INs respectively. (C) Mean song motif similarity and mean IN similarity of pupils to their social tutors and their fathers. Red line represents chance level similarity. * represents p < 0.05, Wilcoxon sign-rank test (D), (E) Mean IN number of pupil vs. mean IN number of father (D) and social tutor (E). Symbols represent individual birds, stars represent birds tutored at McGill, triangles represent birds tutored at IISER Pune. Red line represents regression line. Blue triangle in (C), (D) and (E) represents bird shown in (B).

The tutoring phase lasted ∼1.5 months (phd 34-40 to phd 91-97) for birds tutored at IISER Pune and for 5 days for birds tutored at McGill. Significant song learning is observed for socially tutored birds even after just 5 days of tutoring [35].

#### Isolate birds

5 lab bred male zebra finches (from two different nests) were reared in isolation. After birds had reached nutritional independence, they were separated and kept in visual but not acoustic isolation from other birds. Earlier studies have shown that visual isolation is sufficient to prevent song learning from other birds [36].

### Data analysis

All analyses were conducted using custom written scripts in Matlab (Mathworks).

#### Song segmentation and syllable categorization

Song files were segmented and labeled as described previously [22,23] (see Supplementary Methods for further details). The motif was identified based on the most common sequence of syllables across all bouts. Bouts with one or more motif syllables were considered “song bouts”. Importantly, vocalizations that preceded the first motif of a bout were considered introductory notes (INs). Vocalizations that were also produced outside of song bouts were considered as calls and not as introductory notes. Fig. S1 shows the motif syllables and INs for one example bout for all of the normally reared birds from 16 nests.

#### IN number calculation and associated controls

Before the first motif in each bout, the last set of consecutive INs, with ≤ 500 ms between them, were considered for counting IN number [22,23,37]. For birds with multiple types of INs (30/132 birds had two types of INs, 7/132 birds had 3 or more types of INs), all types of INs were included in the calculation of IN number.

There was variation in the sample size of birds in each nest. To check whether this difference in sample size could influence the correlation in IN number between fathers and normally reared birds, we used a random reassignment procedure to assess significance of the observed correlation. For this procedure, all sons were randomly reassigned to different fathers, while maintaining the actual number of birds per nest, and then the correlation between fathers and sons was calculated. This procedure was repeated 10000 times and the 95% confidence intervals were estimated. This same randomization procedure was also used to assess the significance of the correlation between socially-tutored birds and their tutors. In this case, the total number of socially tutored birds was maintained constant during the random reassignment procedure.

While classifying syllables as INs or motif syllables, we were not blind to the relationship between birds or the experimental group. To control for any potential biases that this might have introduced in our classification of INs and motif syllables, we also used a script based categorization of INs and motif syllables. All song bouts were considered and syllable sequences within a bout with ≤ 500 ms inter-syllable interval were considered. Across all these syllable sequences, any syllable that was repeated (self-transition probability > 0) and occurred as the first syllable of a bout in > 2% of song bouts, was considered an IN. Syllables that were present in 90% of bouts and did not occur as the first syllable of a bout were considered motif syllables. The remaining syllables that did not satisfy either of these criteria were classified as calls.

#### Comparison of IN acoustic structure

Acoustic structure similarity for both motifs and INs was computed using Sound Analysis Pro (http://soundanalysispro.com/) [38]. Twenty randomly picked motifs and 10 randomly picked INs were used for calculating similarity. Asymmetric, time course similarity was calculated for motif to allow for potential differences in syllable order. Symmetric, time course similarity was used for comparing IN acoustic structure as these are individual syllables. To control for this difference in similarity calculation for INs and motifs, we also calculated symmetric, time course similarity for motifs. Irrespective of the mode of similarity calculation for motifs, we found no significant correlation between IN similarity and motif similarity across birds and their tutors confirming independent learning of motifs and INs.

#### Statistics

Pearson correlation coefficient was used for all correlations and was calculated using the matlab function corrcoef. Linear fits to data were calculated using the matlab functions polyfit and polyval. Wilcoxon rank sum test was used for calculating the significance of song and IN similarity for normally reared and playback tutored birds. Wilcoxon sign rank test was used for comparing song and IN similarity of socially tutored pupils with their fathers and their social tutors.

## RESULTS

### IN acoustic structure and mean IN number are learned from a tutor

To examine the degree to which IN acoustic structure and mean IN number are learned, we first compared these properties for birds that had been reared normally with their fathers (n=16 nests, n=65 birds, Fig. 1C, Fig. S1). INs (and song) of fathers were acoustically more similar to INs (and song) of their sons as compared to INs (and song) of unrelated birds (Fig. 1D; p = 0.041 for INs and p < 0.001 for song, Wilcoxon rank sum test). Mean IN number before song was also significantly correlated between fathers and their sons (Fig. 1E). This correlation in mean IN number was not influenced by removal of individual nests from the analysis (Fig. S2A) and was significantly different from the correlations obtained by randomly re-assigning individual birds to different nests (Fig. S2B). Further, this correlation in mean IN number was significant even when syllables in individual birds were categorized into INs or song syllables based on pre-set rules (Fig. S2C, see Methods). These results showed that mean IN number before song and IN acoustic structure were correlated between fathers and their sons.

To further test the contribution of learning to IN repetition and structure, we used a social-tutoring paradigm where juvenile zebra finches were kept in visual and acoustic contact with an adult male different from their father [34–36] (Fig. 2A; n=14 birds; see Methods for details of tutoring). Importantly, both IN acoustic structure and mean IN number for the social tutor were different from those of the juvenile’s biological father (see example in Fig. 2B and Methods). INs of such socially-tutored birds were acoustically more similar to those of their social tutors than those of their fathers (Fig. 2C; p = 0.203 for INs, p < 0.001 for motif, Wilcoxon sign-rank test). Mean IN number before song was also correlated with mean IN number of the social tutor but not of the father (Fig. 2D, 2E). In fact, we observed a negative correlation between mean IN number of socially tutored birds and mean IN number of their fathers (Fig. 2D) reflecting our choice of social tutors with IN number different from their fathers (see Methods). The correlation with social tutor was significantly different from that obtained by randomly re-assigning birds to different social tutors (Fig. S3A) and was significant even when syllables were categorized based on pre-set rules (Fig. S3B). These two sets of results indicate that the number and acoustic structure of INs are learned from a tutor (father for normally reared birds and social tutor for socially-tutored birds).

### Accurate learning of INs is independent of accurate learning of song

Previous studies have suggested that INs represent motor preparation before song [22,23]. If INs represent motor preparation for song, similarity in song between individual birds and their tutors should result in similarity in IN number and/or IN acoustic structure between birds and their tutors. However, across all normally reared and socially tutored birds, the degree of song similarity between birds and their tutors was not correlated with similarity in IN number (Fig. 3A, Fig. S4A) or similarity in IN acoustic structure (Fig. 3B, Fig. S4B) between birds and their tutors (father or social tutor). Further, across all of these birds, mean IN number was not correlated with differences in various temporal (Fig. S5A - duration of song; Fig. S5B – duration of first song syllable) and spectral aspects of song (Fig. S5C – mean frequency of first motif syllable, Fig. S5D – complexity of first motif syllable). These results showed that the number of times an IN was repeated was independent of the song that followed and suggest two independent processes involved in IN and song learning.

**FIGURE 3.**
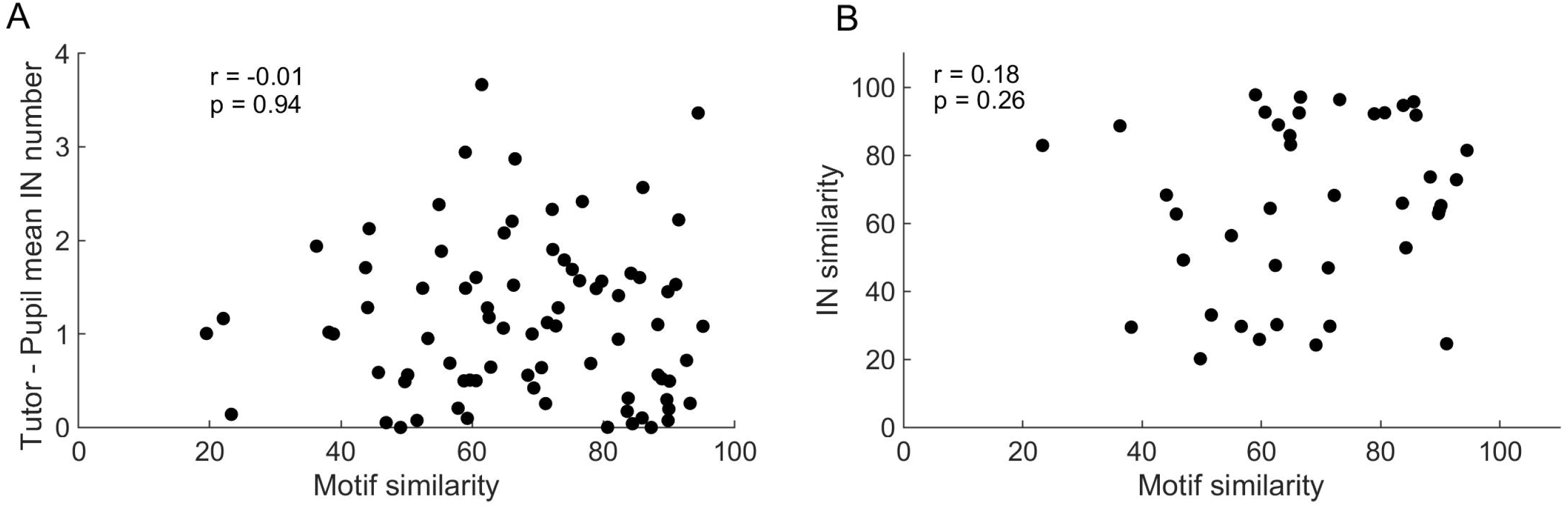
IN number and IN acoustic structure are learned independent of song learning. (A), (B) Motif similarity between tutor (father or social tutor) and pupil vs. absolute difference in mean IN number between tutor and pupil (A) or IN similarity between tutor and pupil (B). Circles represent individual birds. Here motif similarity and IN similarity were calculated using asymmetric, time-course similarity and symmetric, time-course similarity respectively.

### Biological predispositions in IN production and learning

Our results suggest that INs are learned from a tutor just like birds learn their song motifs from their tutor. Previous studies have demonstrated the presence of biological predispositions in zebra finch song learning [30,39]. For example, juvenile zebra finches that are individually tutored with random sequences of syllables converge on similar motif sequences [30]. To identify biological predispositions, if any, in IN production, we experimentally tutored juvenile zebra finches with songs that lacked INs (Fig. 4A, n=22, see Methods for details of tutoring) [30,33,34]. Despite being tutored without INs, juveniles tutored in this manner produced INs before their songs (Fig. 4B for an example). The number of these INs was not correlated those of the father (Fig. 4C). The acoustic similarity of these INs to those of the father was comparable to the similarity with unrelated birds, showing that these INs were acoustically different from those of the father. (Fig. 4D, p = 0.82 for INs and p = 0.71 for motif, Wilcoxon rank sum test). These results suggest more general, species-specific, predispositions in IN production rather than direct genetic components from the father.

**FIGURE 4.**
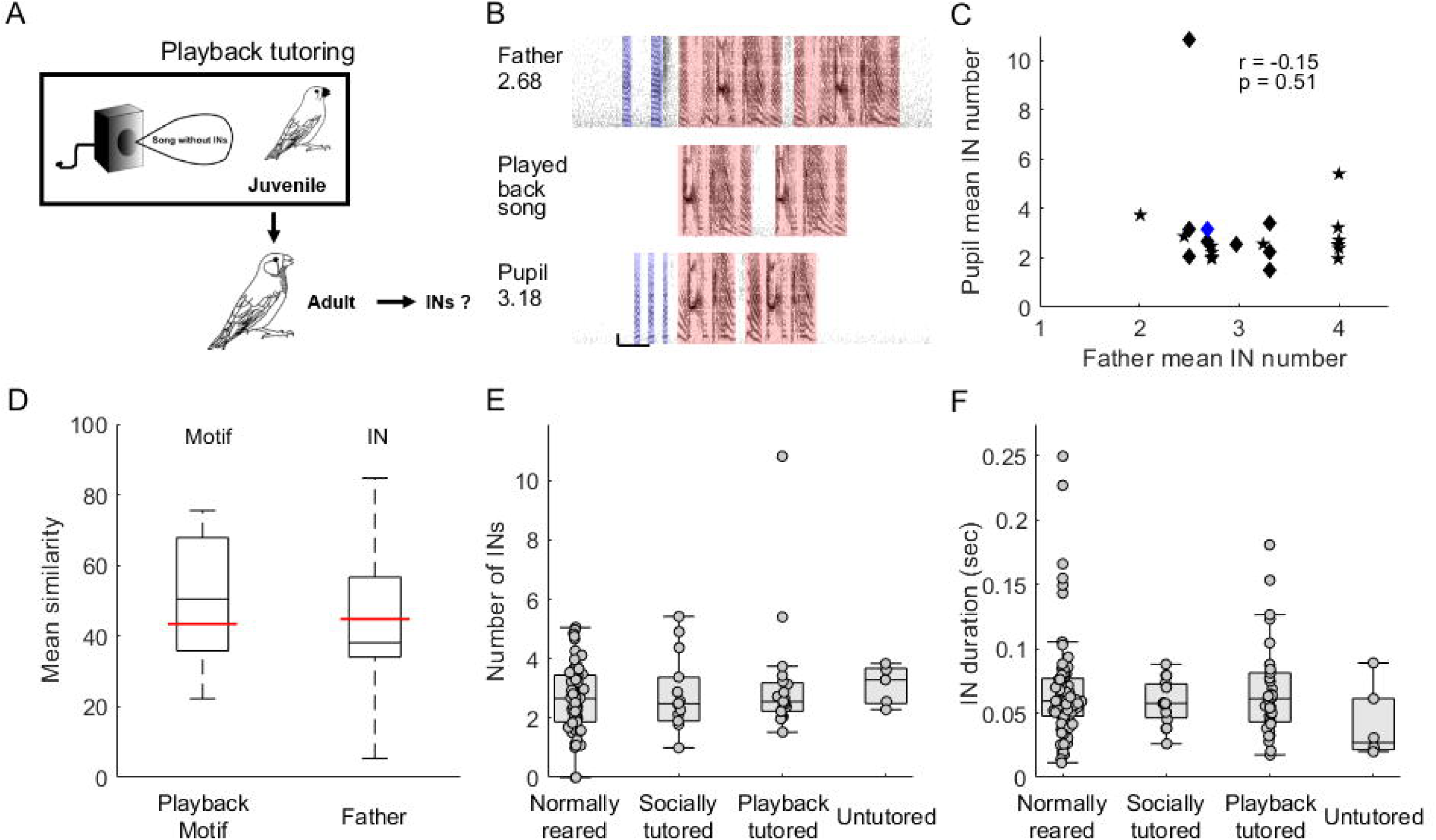
Biological predispositions in IN production. (A) Schematic of playback tutoring paradigm. (B) Example spectrograms for a song bout for the father (top), played back song (middle) and playback-tutored bird (bottom). Blue shaded regions highlight INs and red shaded regions highlight song motifs. Black number on the left of the spectrograms represent mean IN number for that bird. Blue and red numbers represent similarity to father’s INs and played-back motif respectively. (C) Mean IN number for father vs. mean IN number for playback tutored birds. Stars represent birds tutored at McGill, diamonds represent birds tutored at IISER Pune. Blue diamond represents bird shown in (B). (D) Mean motif similarity for playback tutored birds relative to played back motif and mean IN similarity relative to father. Red lines represent random similarity. (E), (F) Box plot representing mean number (E) and duration (F) of INs produced by normally reared (n=65), socially tutored (n=14), playback-tutored (n=22) and untutored birds (n=5). Circles represent INs in individual birds. Multiple IN types in a given bird are represented separately, so the total number of points in each category can exceed the number of birds.

Another way to reveal biological predispositions in IN production is to analyze INs in the songs of untutored birds. As observed in a previous study [10], we found that untutored birds produce INs before their songs (n=5, see Methods for details of untutored birds). Interestingly, the mean number of such INs produced before song (Fig. 4E; p = 0.57, one-way ANOVA) and the duration of these INs (Fig. 4F; p = 0.3662, one-way ANOVA) was similar across all categories of birds, irrespective of tutoring experience. Other features of INs differed between birds experimentally tutored without INs and birds normally reared or socially tutored with INs (Fig. S6A, S6B, S6C) highlighting the role of learning. Thus, on average, normally reared, socially tutored, operantly tutored and untutored birds produced three, 60ms long, INs before starting their songs (Fig. 4E, 4F). This bias to produce ∼3INs before song can also be observed in the data for birds tutored with songs that contained INs. Juveniles tutored by adults that produced, on average, < 3 INs tended to produce more INs than their tutor (Fig. 1E, 2E). On the other hand, juveniles tutored by adults that produce, on average, > 3 INs tended to produce fewer INs than their tutor (Fig. 1E, 2E).

## DISCUSSION

The complex vocal displays of many songbirds begin with the repetition of simple, introductory notes (INs) before the production of their learned song. Here, we show that INs, just like elements in song, are learned and shaped by biological predispositions. Specifically, we show that the mean number of times an IN is repeated and the acoustic structure of INs are learned from a tutor independent of song learning. We also show that birds tutored without INs and untutored birds produce INs with similar duration and number before their songs as birds tutored with INs, revealing the existence of biological predispositions in IN production. Taken together, these results suggest that acquisition of the complex vocal displays of songbirds involves the independent acquisition of INs and song.

### Learning of IN number and structure

How do birds learn the number and structure of INs? Earlier studies have shown the presence of multiple strategies for learning song that involve either learning the sequence first and gradually changing the acoustic structure or learning each syllable sequentially [33,40]. Similarly, juvenile birds could potentially learn IN number first while gradually modifying IN acoustic structure. Alternatively, birds could learn the structure of INs and then gradually learn to transition to the motif after the correct number of INs are repeated. Further studies of IN development in young birds could shed more light on the process of IN learning. A more recent study using experimental tutoring also showed that birds preferentially learn syllable structure at the expense of sequence [41]. This predicts accurate learning of IN structure independent of accurate learning of IN number.

### Functional significance of INs and other such introductory gestures

What is the functional role of INs in zebra finch song? Many different roles have been proposed for introductory gestures including an “alerting” function [5,18,19], a species-specific signal that aids learning of song [20], a “local-dialect” signal [21] and a reflection of motor “preparation” [22,23]. While the degree to which INs in birdsong serve an alerting or identification function remain largely unknown, our data showing learning of IN number and IN acoustic structure suggests that these two aspects may not reflect motor preparation. Rather, learning of these two aspects suggests possible communicative functions of INs, such as in individual, regional or species identification.

However, our data also reveals a biological predisposition in IN production. Specifically, we observed that zebra finches are biased to produce short-duration vocalizations as INs, approximately three times before their songs, regardless of their tutoring experiences during development. Interestingly, INs are also seen before the start of song in suboscine birds [13,15,17] where song is not learned and before advertisement calls in other vertebrates that produce unlearned vocalizations including frogs [4] and toadfish [5]. These data support our findings showing the presence of biological predispositions in IN production even before learned vocalizations, like birdsong.

Overall, irrespective of mechanism and function, our results show that the zebra finch can be an excellent model system to understand how introductory gestures are produced, how they transition to the complex vocal display and their possible function.

## SUPPLEMENTARY INFORMATION

Supplementary information includes supplemental methods and 6 supplemental figures.

## ACKNOWLEDGMENTS

We would like to thank Prakash Raut for bird colony maintenance and Yining Chen for assistance with data collection. We would like to thank Aurnab Ghose, Deepa Subramanyam, Michael Long, Dave Mets, Hamish Mehaffey, Mimi Kao, Anand Krishnan and members of the Rajan and Krishnan labs for useful discussions and comments on earlier versions of the manuscript.

## AUTHOR CONTRIBUTIONS

SK and RR designed the experiments with inputs from JTS. SK and VY carried out the experiments. LS and JTS provided additional data for playback-tutored and socially tutored birds. SK, VY and RR analyzed the data. SK and RR wrote the manuscript in consultation with VY, LS and JTS.

