## Supplemental Methods and Supplemental Figures for "Introductory gestures before songbird vocal displays are shaped by learning and biological predispositions"

### SUPPLEMENTAL INFORMATION

#### SUPPLEMENTAL METHODS

##### Song recordings

Songs were recorded on a computer using either a sound card (44.1 kHz for 128 birds) or a data acquisition board (32kHz for 4 birds; National Instruments, NIDAQ PCI). Sound was either recorded continuously for extended periods of time (continuous recordings) or small chunks of data were recorded when the signal crossed a pre-set threshold (triggered recordings). In triggered recordings, 2-3 seconds of data before and after threshold crossing were saved along with the data that crossed threshold. All birds were recorded when they were adults (> 85 days post-hatch) and only undirected songs in the absence of any other bird were recorded. For most analyses, only files with > 2s of silence before and after song bouts in a file were considered. However for the analysis of songs of 4 fathers, we considered files with > 1.9s of silence before and after, as there were not enough files with >2s silence.

##### Experimental groups

We used 4 different experimental groups in our study.

*Normally reared birds*

Twelve of the fathers were purchased from outside. The four lab-bred birds used as fathers were not included as offspring for their nests to avoid data duplication. For two of the nests, two different females were sequentially used for breeding, and offspring from both females were included as part of the nest. For a small number of birds (5/81), we combined data across multiple days to collect > 35 bouts for analysis (n=3 birds, 2 consecutive days combined; n=1 bird, 3 days, spread across a period of 10 days, combined; n=1 bird, 4 days, spread across a period of 6 days).

*Playback tutored birds*

Tutoring procedures varied slightly across locations. McGill birds were operantly tutored using perch hops. Briefly, each time juveniles hopped on a perch, they were presented with one bout of song from a speaker placed outside the cage. Each day was divided into 3 periods and birds were presented a maximum of 10 song bouts during each period. After the maximum number had been reached during a period, further perch hops did not elicit song playback. Each bout consisted of 4 motifs consisting of the same 5 distinct syllables. Importantly, repetitive INs were not part of the song playback.

Birds at IISER Pune were tutored using a similar paradigm. Briefly, a dummy bird was placed in the cage with the juvenile and each time the juvenile pecked the dummy, songs were played back through a speaker placed outside the cage behind the dummy. For six of the birds, each peck elicited the playback of a song bout consisting of two motifs without INs (see Fig. 6A for example). For three of the birds, each peck elicited either a long call or a song bout with two motifs without INs. Birds were tutored for an hour in the morning and an hour in the evening. There was no limit on the total number of songs or calls played back. If there was a long period (~10-15 minutes) during which the juvenile did not peck the dummy, one song bout was played back passively. For 3 of the birds, stimuli were also passively played back at the end of the session (10 song bouts and 10 bouts of long calls). Tutoring lasted 1-2 months (phd 25-30 to phd 60-93). Outside of the tutoring sessions, playback-tutored birds were kept in visual but not acoustic isolation from other birds. Two birds were tutored together in a cage with a dummy bird.

**Data analysis**

All analyses were conducted using custom written scripts in Matlab (Mathworks).

*Song segmentation and syllable categorization*

Zebra finch songs are produced in bouts that consist of multiple renditions of a motif (a stereotyped sequence of syllables). Song files were first segmented using an amplitude threshold with a minimum syllable duration of 10 ms and minimum inter-syllable gap of 5ms. Syllables were labeled with a modified template matching algorithm [1] and manually checked. Groups of syllables with silence > 2s before and after the group were considered as “bouts”.

*Comparison of IN acoustic structure*

For calculating chance level similarity, mean similarity value between the father of a nest and one randomly selected juvenile from each of the other nests was calculated. For birds with multiple INs, similarity was calculated for each IN type and the average was used to represent IN acoustic structure similarity.

#### SUPPLEMENTAL FIGURE LEGENDS

**FIGURE S1** **Spectrograms of fathers and their sons from all nests**

Each vertical black box includes spectrograms of one song bout for fathers (top, enclosed within a green box) and their sons for one of the 16 nests used in our study (Nest a to Nest p). Blue shading highlights INs and red shading highlights motifs. Calls at the beginning of the bout or between motifs are not highlighted.

**FIGURE S2** **Control analyses validating observed correlation in mean IN number between fathers and their sons**

(A) Circles represent correlation co-efficients obtained by removing one nest at a time from the original data. Diamond represents observed correlation coefficient calculated with all nests. All co-efficients were significant ($p < 0.05$, Pearson's correlation co-efficient). (B) Distribution of correlation coefficient values obtained by randomly re-assigning sons to different nests 10,000 times while maintaining the same number of birds for each nest from the actual data. Blue dashed line represents the correlation coefficient calculated using the actual data. Red dashed lines represent the 95% confidence interval. (C) Mean IN number of father vs. mean IN number of son calculated after using pre-set rules to categorise syllables into INs and motif syllables (see Methods). Grey circles represent individual birds, black squares and whiskers represent mean and s.e.m. for individual nests and red line represents regression line.

**FIGURE S3** **Control analyses validating observed correlation in mean IN number between pupils and their social tutors**

(A) Distribution of correlation coefficient values obtained by randomly re-assigning pupils to different father / social tutor combinations 10,000 times. Correlation between pupil and father (left) and correlation between pupil and social tutor (right). Blue dashed line represents the correlation coefficient calculated using the actual data. Red dashed lines represent the 95% confidence interval. (B) Mean IN number of father vs. mean IN number of pupil (left) and mean IN number of social tutor vs. mean IN number of pupil (right) calculated after using pre-set rules to categorise syllables into INs and motif syllables (see Methods). Symbols represent individual birds tutored in different labs; triangles – IISER Pune, stars – McGill. Red lines represents regression line.

**FIGURE S4** **IN number and IN acoustic structure are learned independently**

(A), (B) Motif similarity between tutor (father or social tutor) and pupil vs. absolute difference in mean IN number between tutor and pupil (A) or IN similarity between tutor and pupil (B). Circles represent individual birds. Here motif similarity and IN similarity were both calculated using symmetric, time-course similarity.

**FIGURE S5 Variation in IN number is not correlated with variation in song**

(A), (B), (C) and (D) Mean IN number across birds vs. first motif duration (A), first motif syllable duration (B), mean entropy variance of entire motif (C) and mean entropy variance of first motif syllable (D). Circles in all plots represent individual birds. Only normally reared and socially tutored birds are plotted here.

**FIGURE S6 Differences in acoustic properties of INs across zebra finches with different tutoring experiences**

(A), (B), (C) Box plot representing entropy (A), mean frequency (B) and frequency modulation (C) of INs produced by normally reared (n=65), socially tutored (n=14), playback-tutored (n=22) and untutored birds (n=5). Circles represent INs in individual birds. Multiple IN types in a given bird are represented separately, so the total number of points in each category can exceed the number of birds. * represents $p < 0.05$ and *** represents $p < 0.001$ for the post-hoc Tukey-Kramer test after one-way ANOVA. One-way ANOVA p-values are 0.00014 (for entropy), 0.029 (for mean frequency) and 0.058 (for frequency modulation).

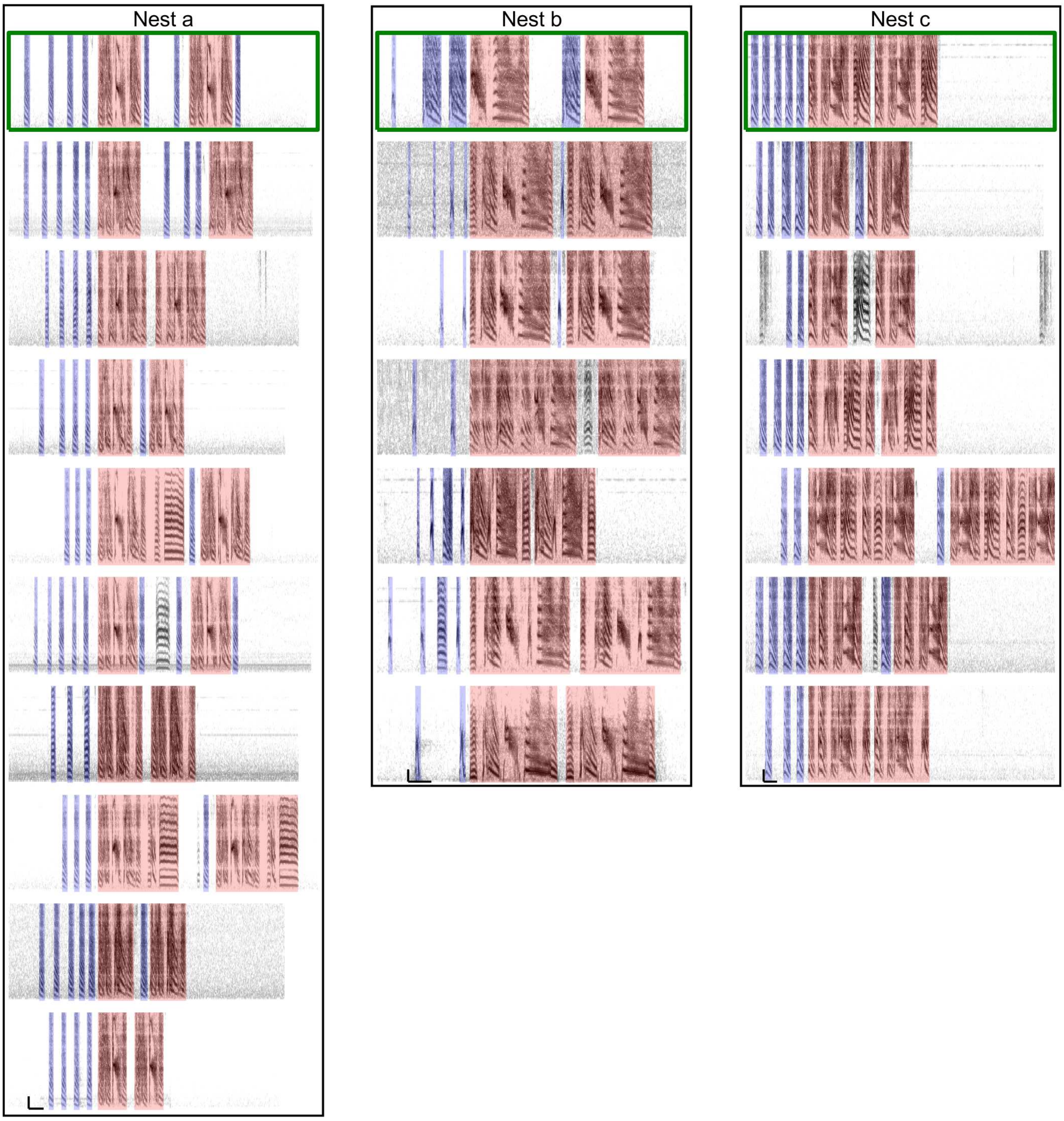


**FIGURE S1** Spectrograms of fathers and their sons from all nests

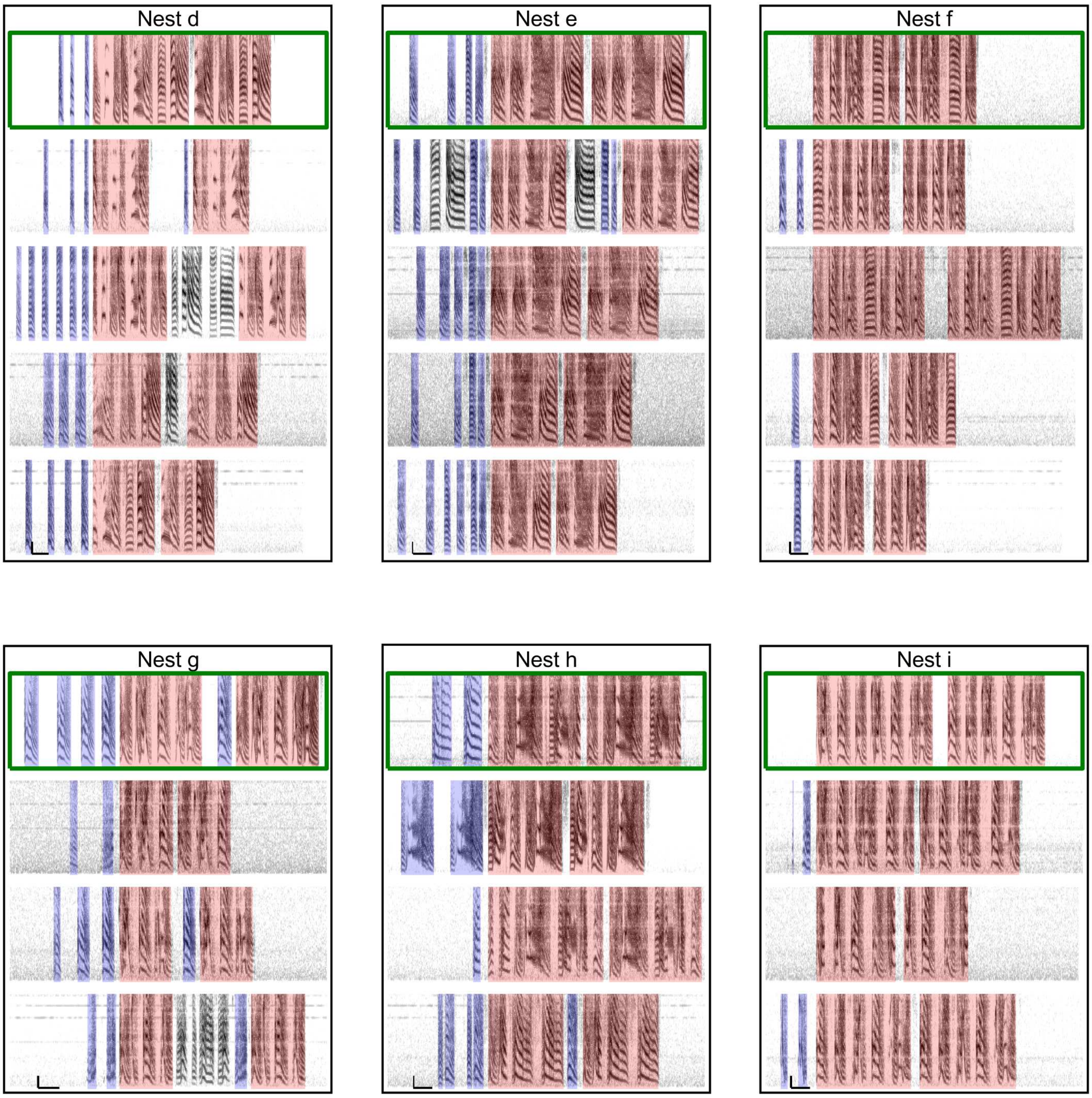


**FIGURE S1** <u>**Spectrograms of fathers and their sons from all nests**</u>

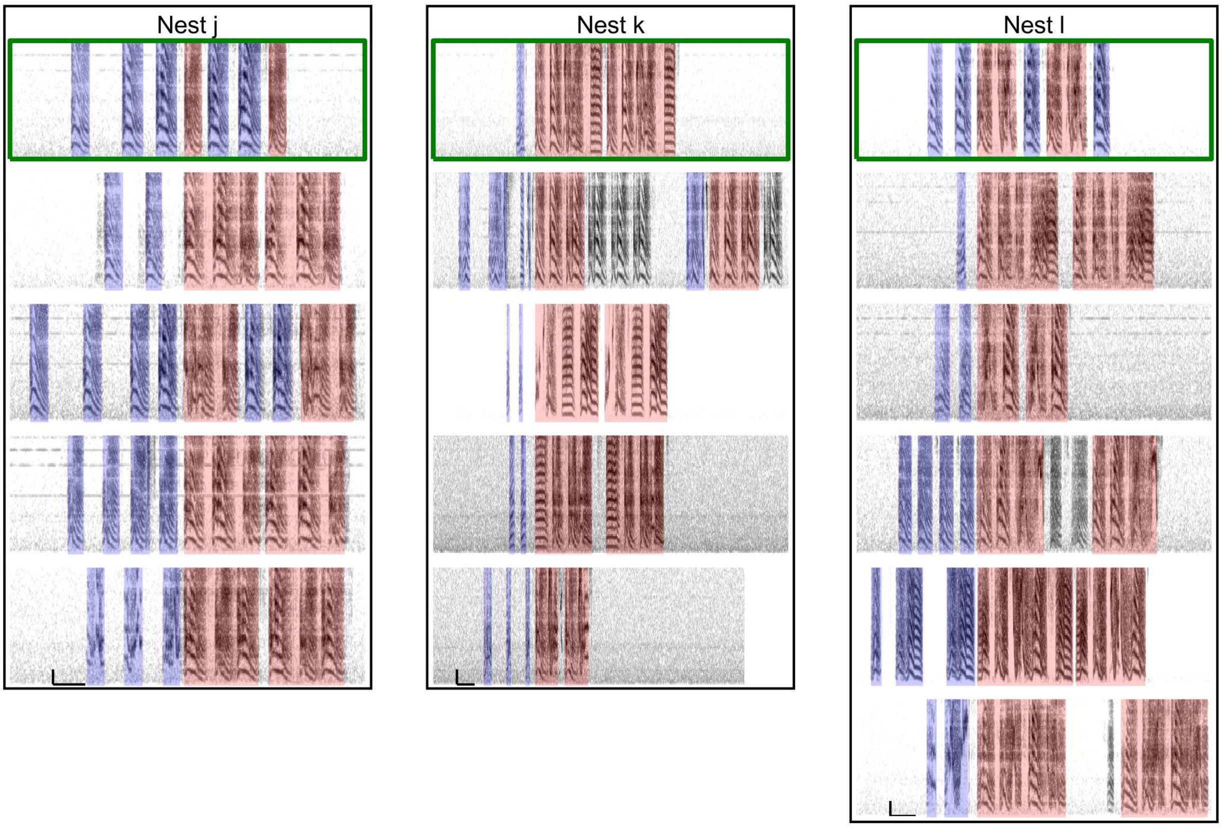


**FIGURE S1** **Spectrograms of fathers and their sons from all nests**

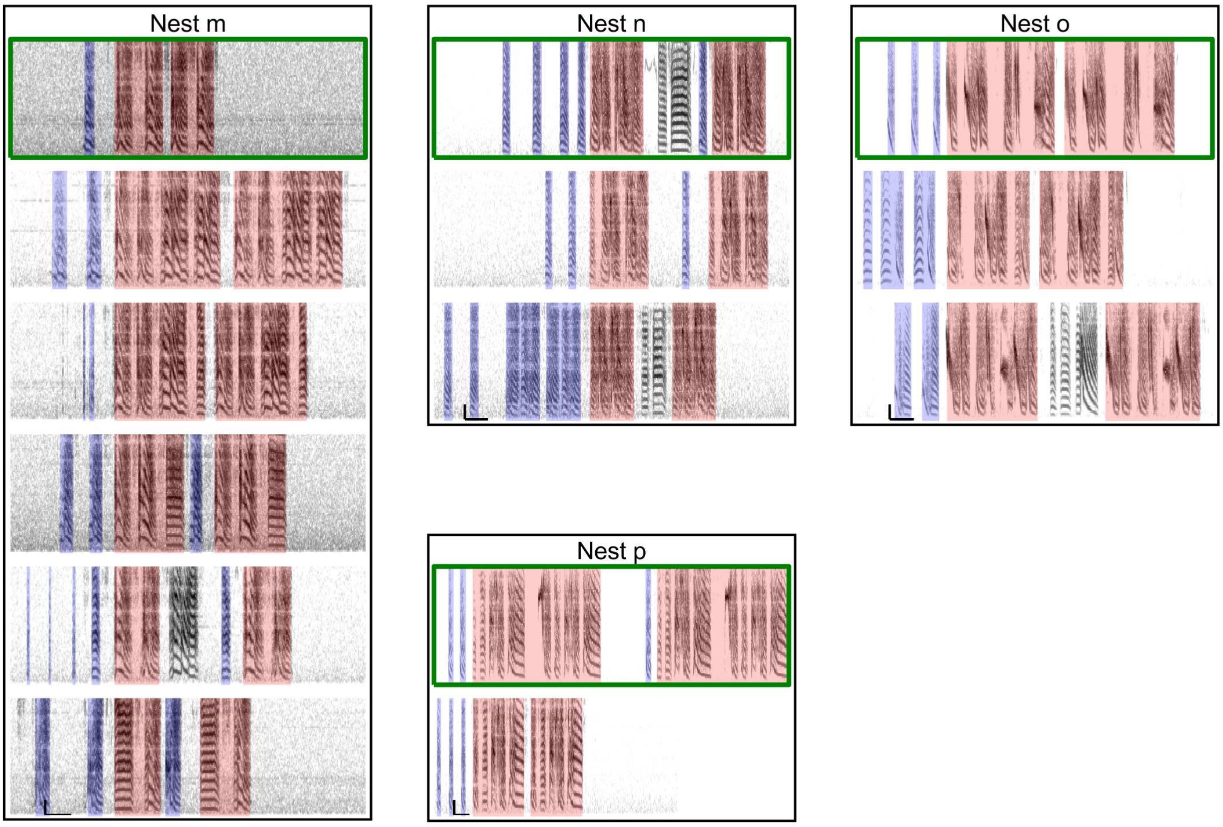


**FIGURE S1** **<u>Spectrograms of fathers and their sons from all nests</u>**

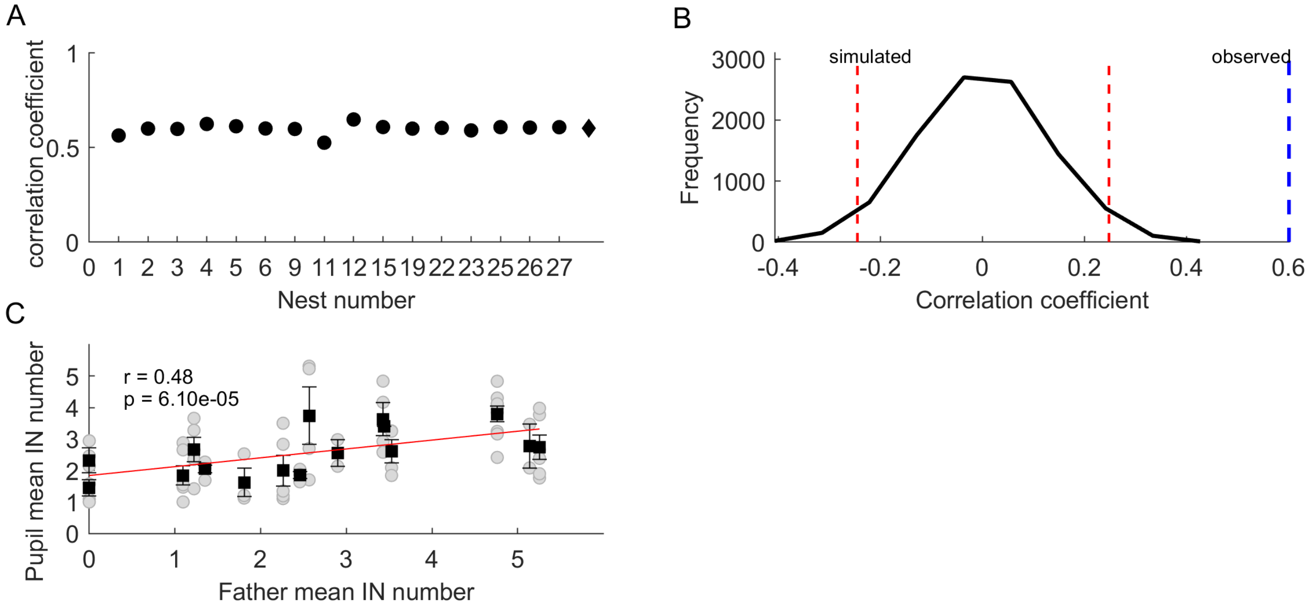


**FIGURE S2** **Control analyses validating observed correlation in mean IN number between fathers and their sons**

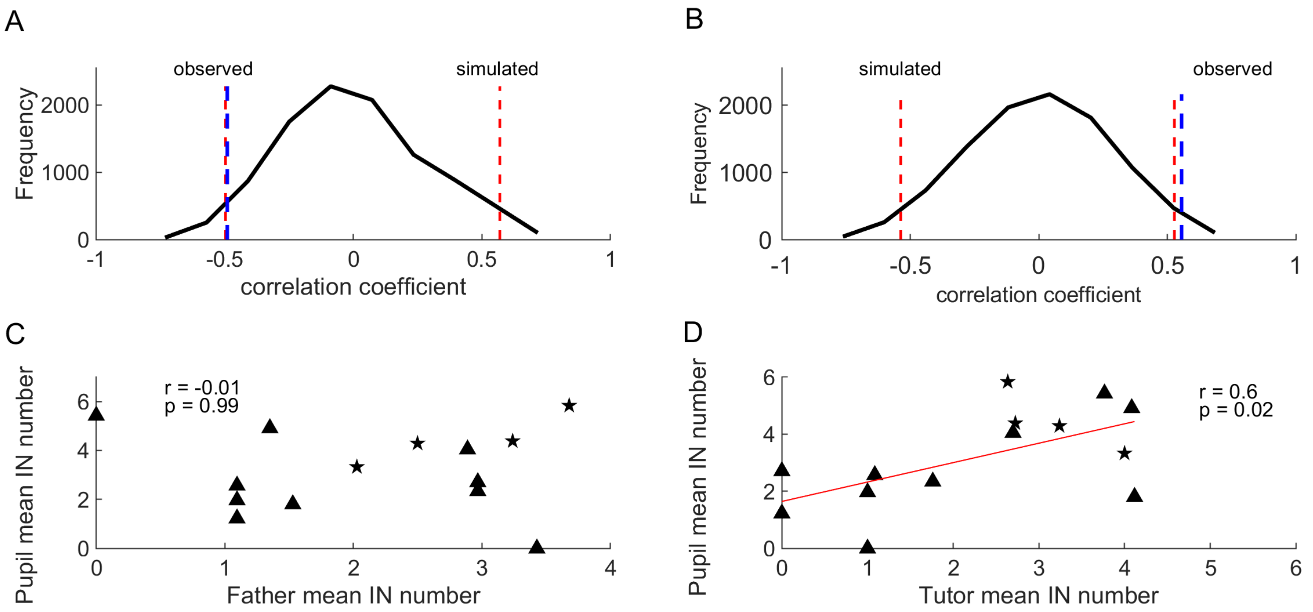


**FIGURE S3** **<u>Control analyses validating observed correlation in mean IN number between pupils and their social tutors</u>**

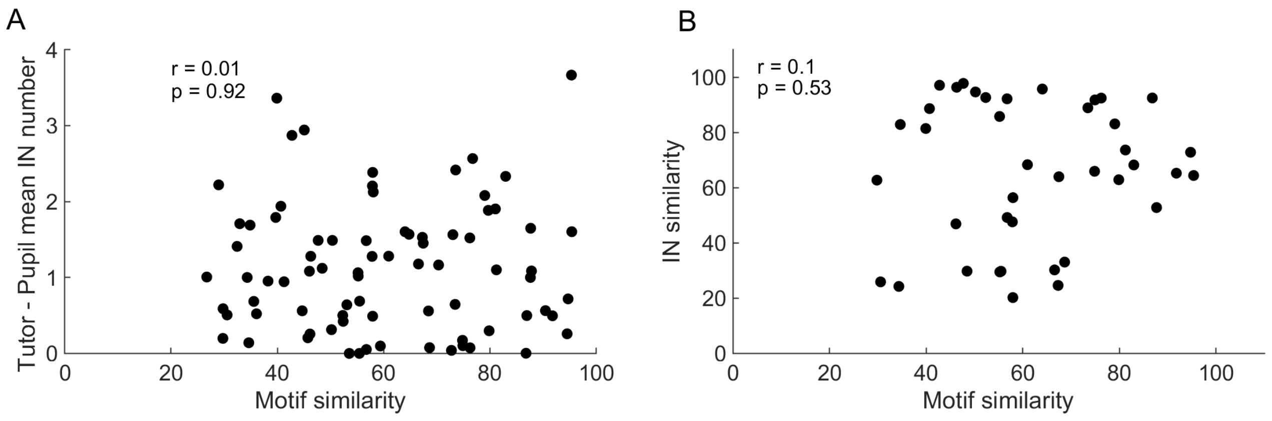


**FIGURE S4** **<u>IN number and IN acoustic structure are learned independently</u>**

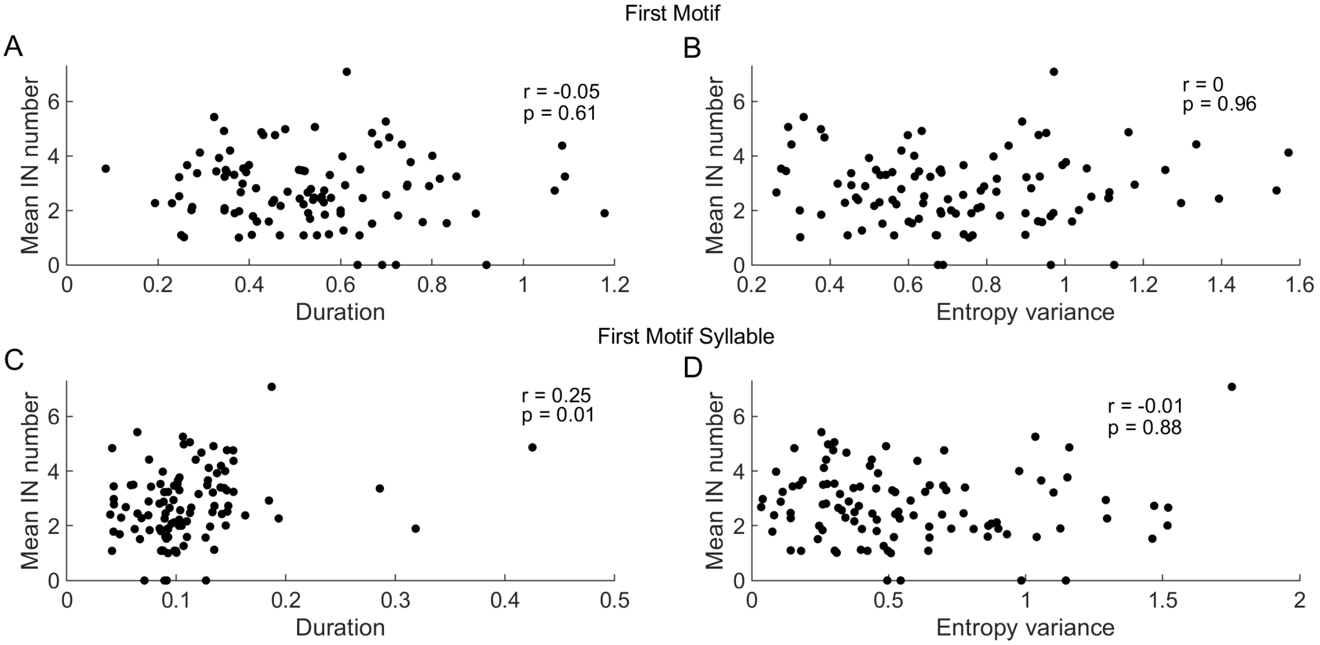


**FIGURE S5** **<u>Variation in IN number is not correlated with variation in song</u>**

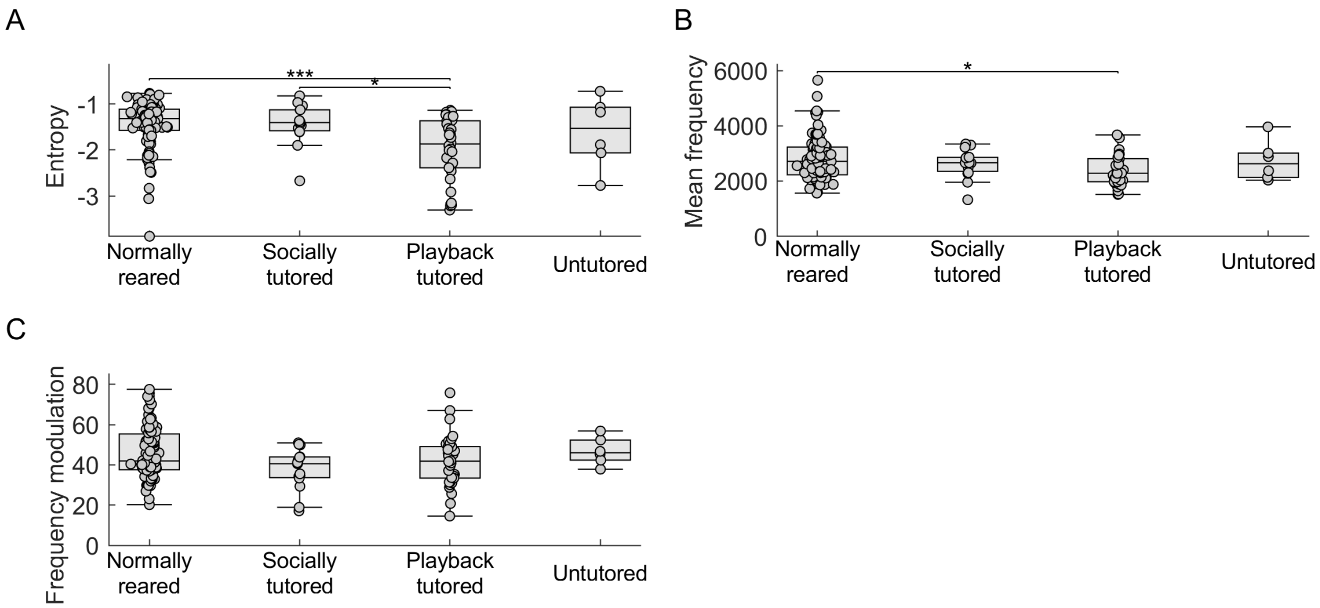


**FIGURE S6 Differences in acoustic properties of INs across zebra finches with different tutoring experiences**
